# Altered basal forebrain regulation of intrinsic brain networks in depressive and anxiety disorders

**DOI:** 10.1101/2024.11.06.622349

**Authors:** Alec J. Jamieson, Trevor Steward, Christopher G Davey, Sevil Ince, James A Agathos, Bradford A Moffat, Rebecca K Glarin, Kim L Felmingham, Ben J Harrison

## Abstract

Depressive and anxiety disorders are characterized by altered connectivity within and between the default mode (DMN) and salience networks. Basal forebrain subdivisions, critical for regulating network activity, remain understudied across these conditions. To address this gap, we analyzed 7-Tesla resting-state functional magnetic resonance imaging data from a transdiagnostic sample (*n* = 70), primarily with depressive and anxiety disorders, and healthy controls (*n* = 77). We used spectral dynamic causal modelling to assess effective connectivity between the medial septum/diagonal band (Ch1-3), nucleus basalis of Meynert (Ch4), ventral pallidum, and DMN and salience networks. Healthy participants showed excitatory connectivity from Ch1-3 to the DMN and from Ch4 to the anterior insula. In contrast, clinical participants exhibited greater inhibitory Ch4 to DMN connectivity and increased excitatory connectivity from Ch4 to the anterior insula. Wide-spread Ch4 connectivity dysfunction may implicate the cholinergic system as a mechanistic and therapeutic target for depressive and anxiety disorders.

## Introduction

Widespread alterations to intrinsic brain networks have been noted across depressive and anxiety disorders^1–3^, particularly in the default mode (DMN) and salience networks^4,5^. Resting-state functional magnetic resonance imaging (fMRI) is commonly used to study these brain networks by measuring spontaneous changes in neural activity while participants are not engaged in any specific task. The relationship between disparate brain regions is typically characterised in terms of functional connectivity, the correlation or similarity between brain signals arising from such regions. Abnormal DMN functional connectivity in major depressive disorder (MDD) has been associated with higher levels of rumination and an increased duration of depressive episodes^6^. Similarly, altered resting-state connectivity, both within the DMN and between the DMN and salience network, has been observed across both social and generalized anxiety disorders^7^ and associated with anxiety symptom severity^8^. Abnormal functioning of these networks has been hypothesized to relate to common symptoms across both conditions. For example, DMN alterations are suggested to contribute to abnormal self-referential processing and rumination, whereas dysfunction in the salience network may relate to abnormal processing of negative stimuli^9–11^. Despite the possible transdiagnostic nature of this brain network dysfunction, reliance on investigations targeting specific categorical diagnoses has limited direct evidence for this relationship^12,13^.

Although the DMN and salience networks are primarily identified as cortical in nature, there is emerging evidence that their function is heavily influenced by subcortical modulatory structures, including the basal forebrain^14–16^. The basal forebrain’s influence over the DMN has been hypothesized to reflect a mechanism for switching between internally and externally directed attention states^15,17^. Activation of cholinergic neurons in the basal forebrain of rodents has been shown to decrease DMN functional connectivity, highlighting a potential role of the basal forebrain in regulating the activity of this network^18^. Notably, the cholinergic component of the basal forebrain is comprised of distinct subdivisions, with heterogeneity illustrated in both their histochemistry and function^19^. These subregions include the medial septal nucleus (cholinergic group 1; Ch1), vertical nucleus of the diagonal band (Ch2), horizontal nucleus of the diagonal band (Ch3), and nucleus basalis of Meynert (Ch4)^19^. Notably, Ch4 demonstrates the highest concentration of cholinergic cells, comprising approximately 90% of its neurons, and represents the major source of cholinergic innervation for the cortical mantle^19^. These basal forebrain subdivisions are largely replicated in resting-state fMRI functional connectivity studies, with Fritz and colleagues illustrating two functional subcomponents congruent with Ch1-3 and Ch4^20^. Importantly these subregions demonstrate differential connectivity patterns with intrinsic cerebral networks, with Ch1-3 exhibiting strong functional connectivity with the anterior DMN, whereas Ch4 activity is correlated with the functioning of the salience and central executive networks^20–22^. The ventral pallidum (VP), a basal forebrain subregion which is predominantly GABAergic^23^, has also demonstrated evidence for controlling transitions between internally and externally guided behavior by regulating DMN activity^14,24,25^. This indicates that different subregions of the basal forebrain may have both distinct and complementary roles in coordinating the activity of different intrinsic brain networks.

The basal forebrain is implicated in functions that are known to be altered across depression and anxiety, including the modulation of circadian rhythms, appetite, motivation, and cognitive flexibility^26–28^. An association between altered central acetylcholine levels and changes in mood and anxiety symptoms have long been noted within the literature^29,30^. Specifically, this research suggests that reductions in acetylcholinesterase activity and increasing acetylcholine levels can induce depression and anxiety symptoms in humans and rodents^31,32^. Rodent work suggests that both lesions in cholinergic neurons and stress-induced hyperactivity of VP GABAergic neurons can produce depression-like behaviors^33,34^. The VP also contains a collection of cholinergic neurons that project to the prefrontal cortex and basolateral amygdala, which may play a key role in valence processing^23,35^. Importantly, these behaviors induced by cholinergic lesions appear reversable through antidepressant treatment, including both ketamine and fluoxetine^33^. Together this indicates that basal forebrain disturbances may be present across anxiety and depressive disorders and contribute to changes observed in the functioning of intrinsic brain networks. However, research on the functioning of the basal forebrain and its subregions remains limited in humans due to the technical limitations of conventional (3-Tesla) fMRI approaches^36^.

To address this gap, we examined the role of basal forebrain subregions, specifically Ch1-3, Ch4, and VP, in coordinating the activity of regions of the DMN and salience network using ultra-high field (7-Tesla) fMRI. Using spectral dynamic causal modeling (DCM)^37,38^ we first characterized the resting-state effective connectivity (i.e., directional influences among brain areas) of these three basal forebrain regions in a sample of healthy controls. Secondly, we examined alterations present in a transdiagnostic clinical sample with depresse and anxiety symptoms compared with this control group. Third, we examined whether depressive and anxiety symptom severity in the clinical group also related to basal forebrain connectivity alterations. The use of DCM allows for inferences on directional interactions through a process of model inversion – identifying connectivity parameters which offer the best trade-off between accuracy (fit of the predicted timeseries to the observed blood oxygen level-dependent (BOLD) signal data) and complexity (deviations from the default model priors), and model inference - comparing the evidence for different models of network architecture^39^. We hypothesized that our clinical sample would be characterized by greater influence from basal forebrain regions to the DMN, particularly from Ch1-3 and the VP due to their greater association with DMN activity^14,20,24,25^. We additionally hypothesized that these same alterations in effective connectivity would be associated with both depressive and anxiety symptom severity in the clinical sample.

## Results

### Demographic and Clinical Differences

As expected, clinical participants’ scores were significantly greater than healthy controls on the total Depression and Anxiety Stress Scale short-form (DASS) and across all three subscales (Table 1). The proportion of female clinical participants was significantly higher than in the healthy control group (X^2^ = 12.26, *p* = .008), consistent with epidemiological estimates^40,41^. However, there were no significant differences in DASS Depression or Anxiety scores between male and female participants within the healthy control (*t*(75) = -1.37, *p* = .174; *t*(75) = .31, *p* = .760, respectively) or clinical groups (*t*(64) = -1.09, *p* = .280; *t*(64) = 1.14, *p* = .257, respectively). No other demographic differences were observed between the two groups (Table 1).

**Table 1.**
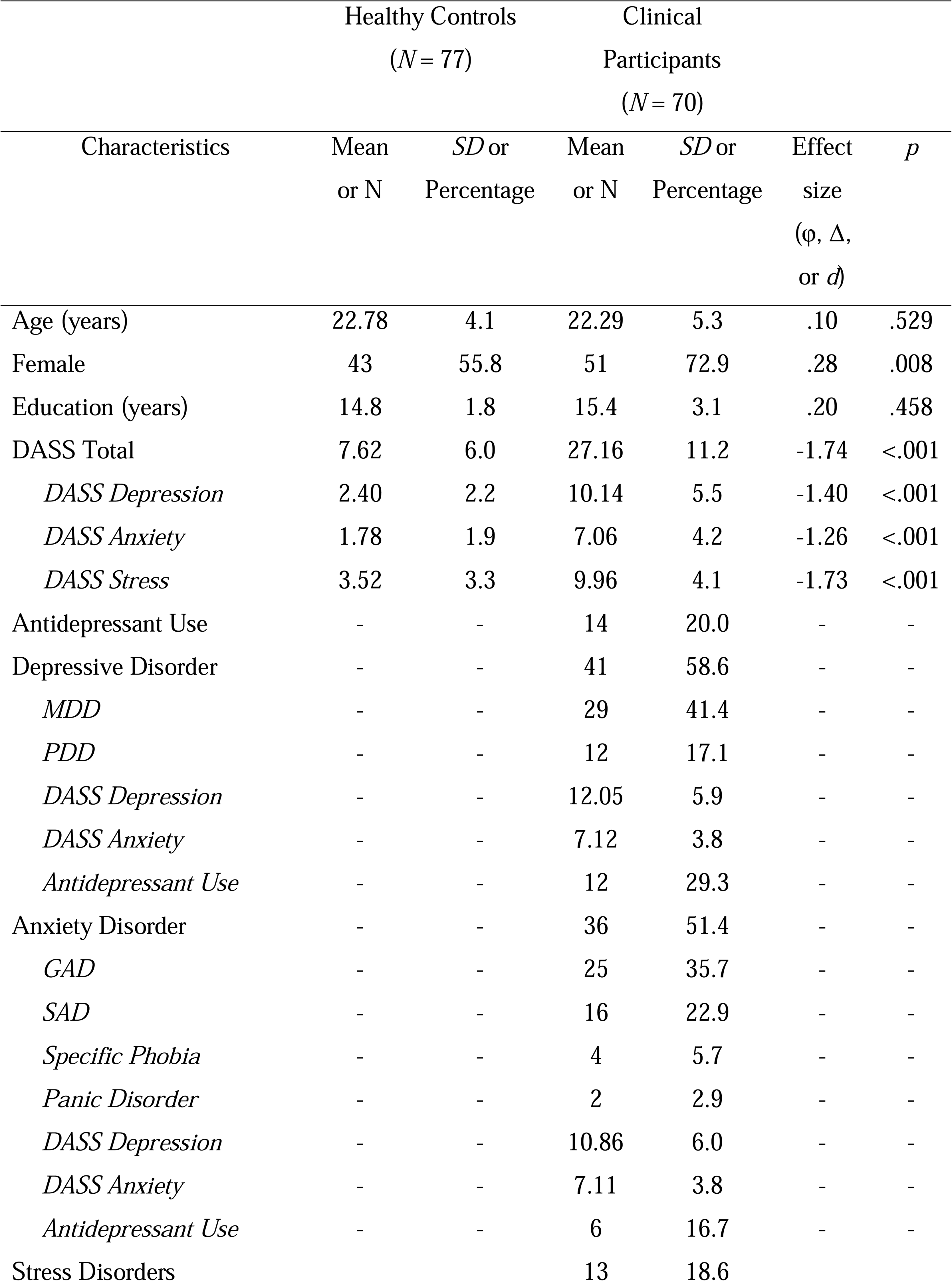
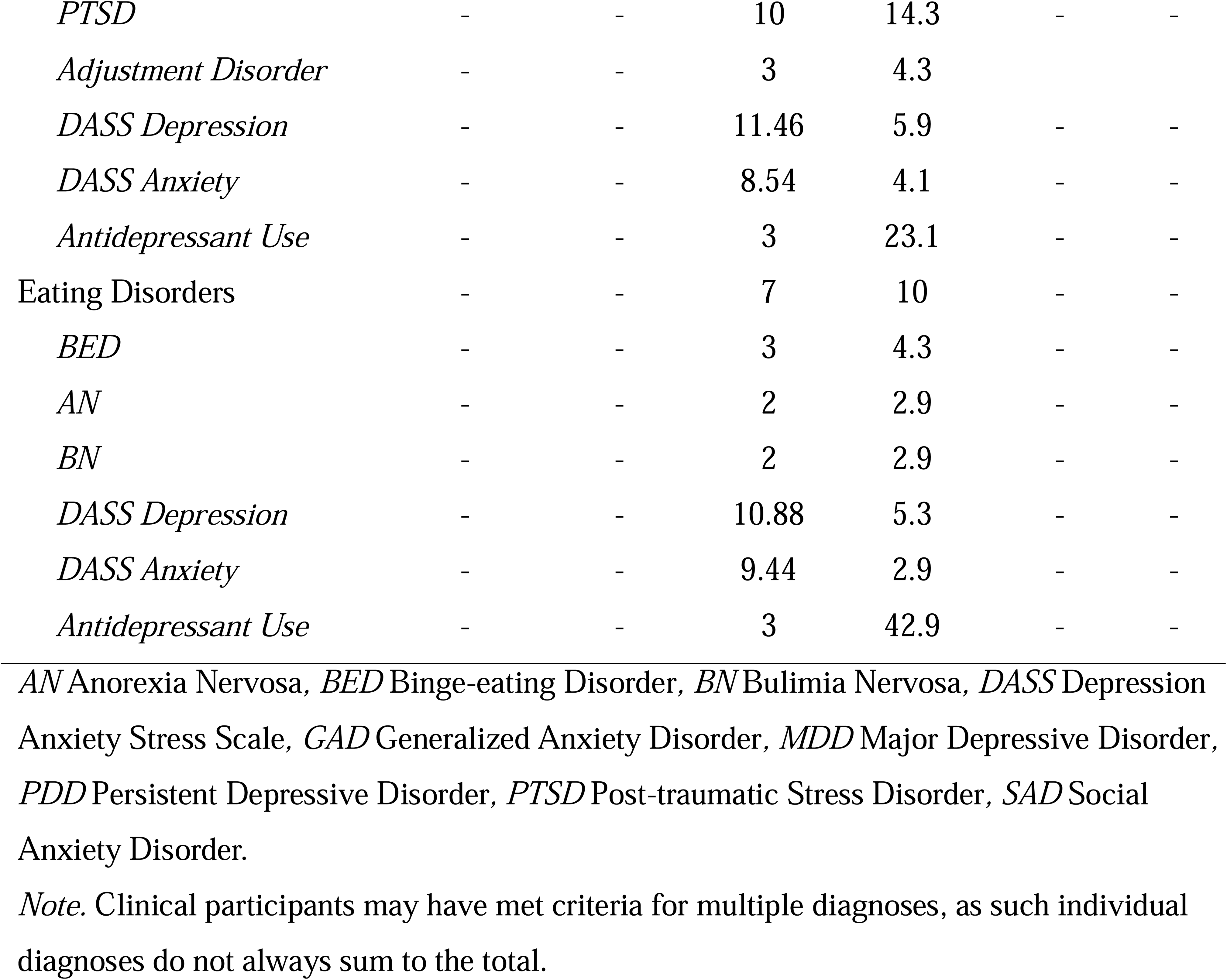
Comparison of Characteristics Between Healthy Controls and Clinical Participants.

| Characteristics | Healthy Controls<br>(*N* = 77) | | Clinical<br>Participants<br>(*N* = 70) | | Effect<br>size<br>($\phi$, $\Delta$,<br>or *d*) | *p* |
| --- | --- | --- | --- | --- | --- | --- |
|  | Mean<br>or *N* | *SD* or<br>Percentage | Mean<br>or *N* | *SD* or<br>Percentage |  |  |
| Age (years) | 22.78 | 4.1 | 22.29 | 5.3 | .10 | .529 |
| Female | 43 | 55.8 | 51 | 72.9 | .28 | .008 |
| Education (years) | 14.8 | 1.8 | 15.4 | 3.1 | .20 | .458 |
| DASS Total | 7.62 | 6.0 | 27.16 | 11.2 | -1.74 | <.001 |
| *DASS Depression* | 2.40 | 2.2 | 10.14 | 5.5 | -1.40 | <.001 |
| *DASS Anxiety* | 1.78 | 1.9 | 7.06 | 4.2 | -1.26 | <.001 |
| *DASS Stress* | 3.52 | 3.3 | 9.96 | 4.1 | -1.73 | <.001 |
| Antidepressant Use | - | - | 14 | 20.0 | - | - |
| Depressive Disorder | - | - | 41 | 58.6 | - | - |
| *MDD* | - | - | 29 | 41.4 | - | - |
| *PDD* | - | - | 12 | 17.1 | - | - |
| *DASS Depression* | - | - | 12.05 | 5.9 | - | - |
| *DASS Anxiety* | - | - | 7.12 | 3.8 | - | - |
| *Antidepressant Use* | - | - | 12 | 29.3 | - | - |
| Anxiety Disorder | - | - | 36 | 51.4 | - | - |
| *GAD* | - | - | 25 | 35.7 | - | - |
| *SAD* | - | - | 16 | 22.9 | - | - |
| *Specific Phobia* | - | - | 4 | 5.7 | - | - |
| *Panic Disorder* | - | - | 2 | 2.9 | - | - |
| *DASS Depression* | - | - | 10.86 | 6.0 | - | - |
| *DASS Anxiety* | - | - | 7.11 | 3.8 | - | - |
| *Antidepressant Use* | - | - | 6 | 16.7 | - | - |
| Stress Disorders |  |  | 13 | 18.6 |  |  |
| *PTSD* | - | - | 10 | 14.3 | - | - |
| *Adjustment Disorder* | - | - | 3 | 4.3 |  |  |
| *DASS Depression* | - | - | 11.46 | 5.9 | - | - |
| *DASS Anxiety* | - | - | 8.54 | 4.1 | - | - |
| *Antidepressant Use* | - | - | 3 | 23.1 | - | - |
| Eating Disorders | - | - | 7 | 10 | - | - |
| *BED* | - | - | 3 | 4.3 | - | - |
| *AN* | - | - | 2 | 2.9 | - | - |
| *BN* | - | - | 2 | 2.9 | - | - |
| *DASS Depression* | - | - | 10.88 | 5.3 | - | - |
| *DASS Anxiety* | - | - | 9.44 | 2.9 | - | - |
| *Antidepressant Use* | - | - | 3 | 42.9 | - | - |

*AN* Anorexia Nervosa, *BED* Binge-eating Disorder, *BN* Bulimia Nervosa, *DASS* Depression Anxiety Stress Scale, *GAD* Generalized Anxiety Disorder, *MDD* Major Depressive Disorder, *PDD* Persistent Depressive Disorder, *PTSD* Post-traumatic Stress Disorder, *SAD* Social Anxiety Disorder.
*Note.* Clinical participants may have met criteria for multiple diagnoses, as such individual diagnoses do not always sum to the total.

### Overview of Effective Connectivity and Model Fit

DCM models the rate of change in activity of a given region as a function of the activity of another region, thus the effect size of this connectivity is measured in hertz (Hz). Positive values are indicative of putative excitation between regions while negative values represent putative inhibition. Heuristics for determining the relative size of these effects can be found in Supplementary Methods and Supplementary Table 1. Self or within-region connections are conversely represented by unitless log scaling parameters that modify the default inhibitory connectivity of -0.5 Hz. As such, positive values for these parameters are indicative of greater inhibition whereas negative values represent reduced inhibition. Based on our aim to examine effective connectivity of Ch1-3, Ch4, and VP with regions of the DMN and salience network, we constructed a DCM network comprised of these basal forebrain regions in addition to the ventromedial prefrontal cortex (vmPFC), posterior cingulate cortex (PCC), and left and right inferior parietal lobules (IPL), as well as the dorsal anterior cingulate (dACC) and left and right anterior insula (*Fig. 1*). For the second-level Parametric Empirical Bayes (PEB) model, a posterior probability (PP) threshold of greater than 0.99 was used to identify those parameters which had a *very strong* amount of evidence^39^.

**Fig. 1.**
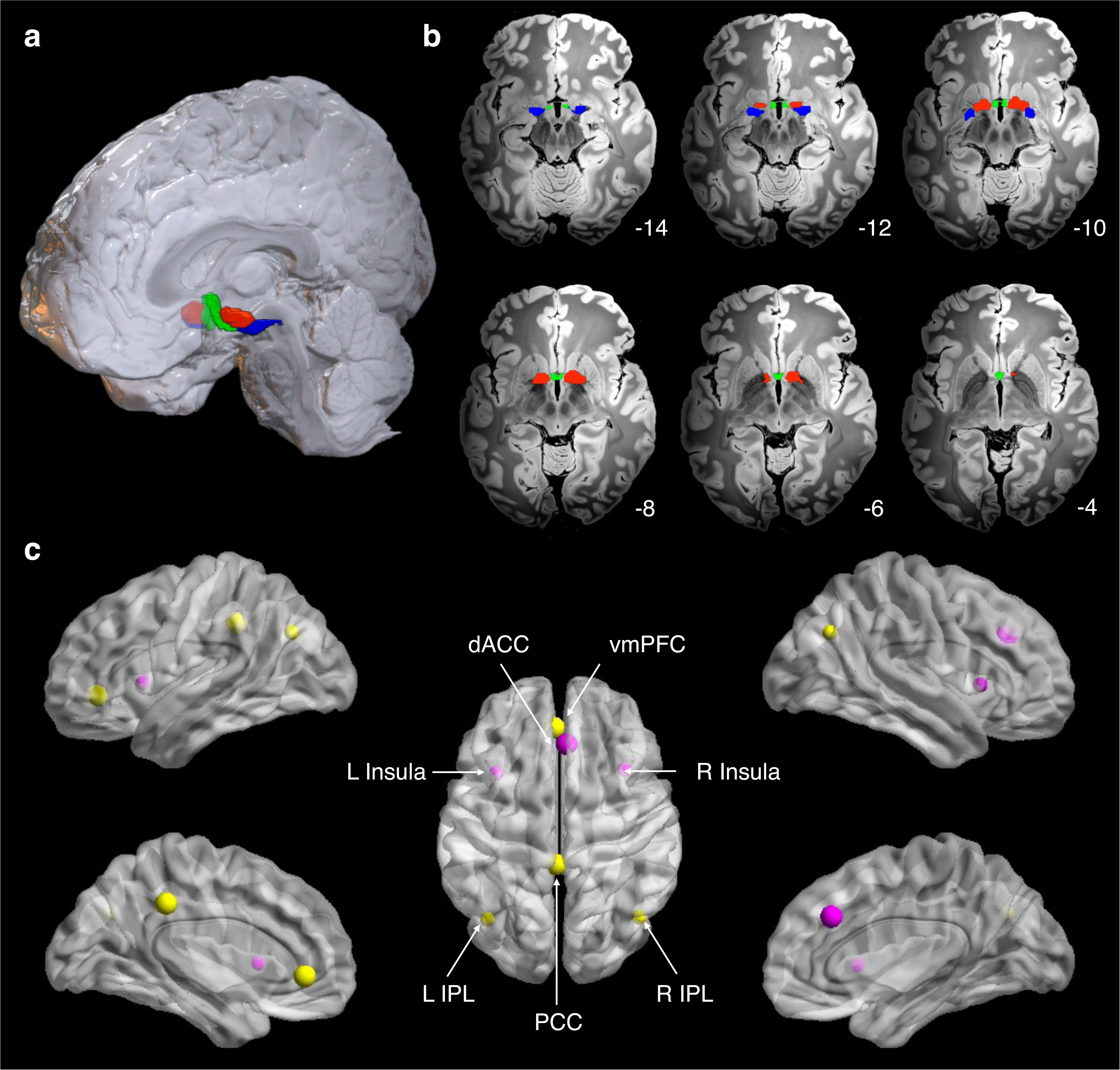
Visualization of the brain regions used in the spectral dynamic causal modelling analysis. **a**, Render and **b**, axial slices of the basal forebrain regions used, Ch1-3 (green), Ch4 (blue), and VP (red) masks on the “Synthesized_FLASH25” (500 μm, MNI space) ex vivo template^101^. Numbers indicate position o0f individual sluices along the z-axis (from inferior to superior). Ch1-3 and Ch4 masks were derived from Zaborszky and colleagues^88^, whereas the VP mask was sourced from Pauli and colleagues^89^. **c**, Visualization of cortical brain regions including default mode network regions (yellow) and salience network regions (pink), weighted by the size of their extracted regions of interest. Render visualized using BrainNet Viewer^102^. Left = Left. *Ch1-3* medial septal nucleus, *Ch4* nucleus basalis of Meynert*, dACC* dorsal anterior cingulate cortex, *IPL* inferior partial lobule, *L* left, *PCC* posterior cingulate cortex, *R* right, *vmPFC* ventromedial prefrontal cortex, VP ventral pallidum.

The coefficient of determination (R^2^) was calculated (using *spm_dcm_fmri_check*) to assess the fit of our DCM model. This reflects the proportion of variance in the observed data which is explained by the model, with values closer to 1 reflecting better model fit. This revealed an average explained variance of 87.3% ± 4% across all subjects, indicating that the specified model was capturing a large degree of the observed changes in the BOLD signal over time.

### Effective Connectivity in Healthy Controls

To provide an overview of the typical functioning of this circuitry, we first characterized the average connectivity pattern illustrated in healthy controls (*Fig. 2A*). Consistent with effects identified in the functional connectivity analysis (*Supplementary Fig. S1*), healthy controls demonstrated an excitatory influence from Ch1-3 to regions of the DMN and an inhibitory influence on regions of the salience network, whereas this relationship was inverted for Ch4 and VP.

**Fig. 2.**
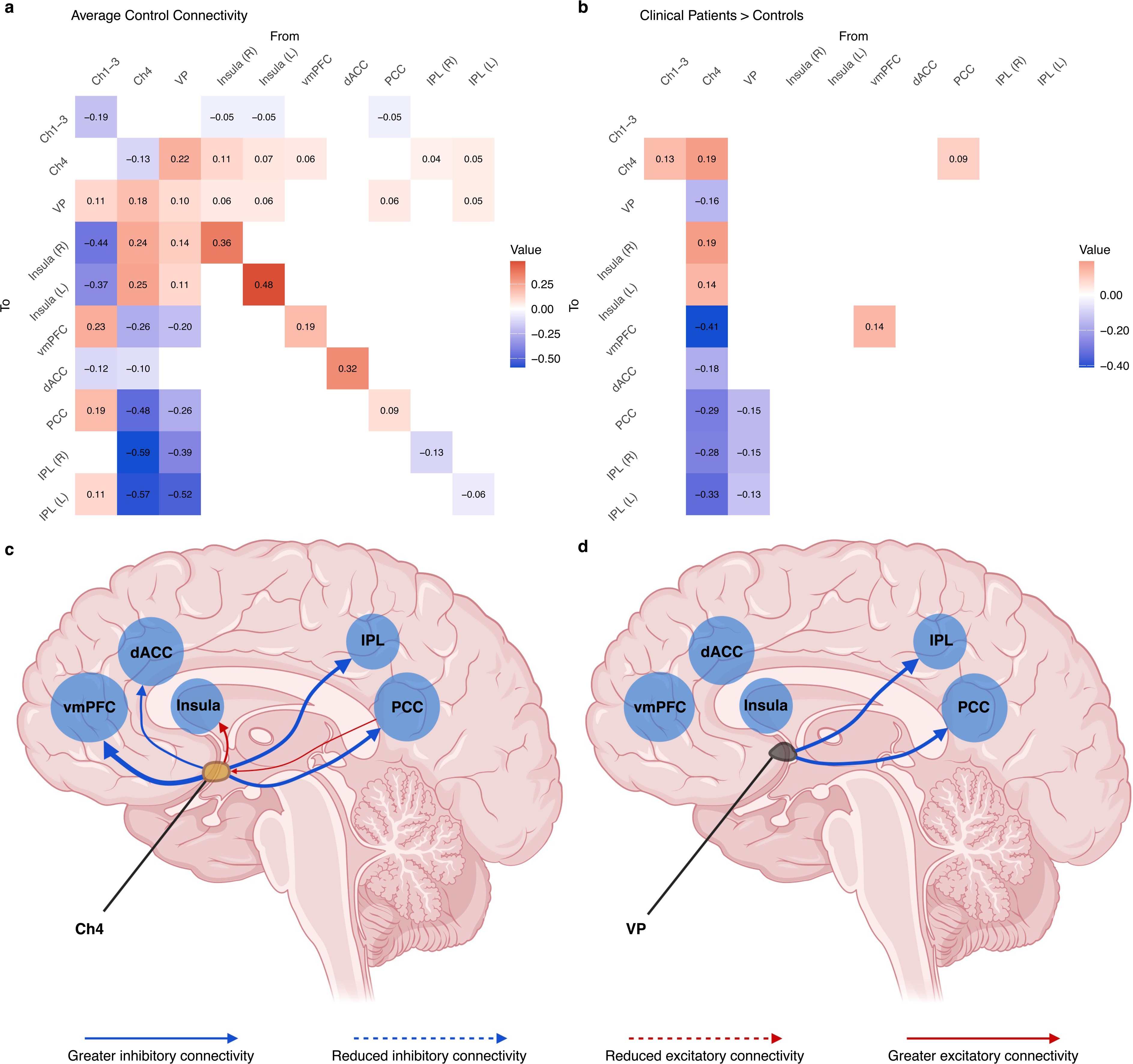
Effective connectivity of the basal forebrain across healthy controls and clinical participants. Adjacency matrices show the average effects observed in **a**, healthy controls and **b**, the difference between clinical participants and healthy controls. For **a**, red cells indicate excitatory connections whereas, blue cells indicate inhibitory connections. For **b**, these values add (red) to or subtract (blue) from the connectivity in **a**,. Cells representing connectivity between two regions are measured in Hz and diagonal cells indicate inhibitory self-connections which are unitless log scaling parameters. **c**, and **d**, illustrate differences in effective connectivity from the basal forebrain to cortical brain regions in clinical participants compared with healthy controls. Results indicate effects involving C) Ch4 (yellow) and D) VP (black). Arrows have been weighted to indicate the relative size of the effects. Image created with BioRender (www.biorender.com). *Ch1-3* medial septal nucleus, *Ch4* nucleus basalis of Meynert*, dACC* dorsal anterior cingulate cortex, *IPL* inferior partial lobule, *PCC* posterior cingulate cortex, *vmPFC* ventromedial prefrontal cortex, VP ventral pallidum.

Specifically, Ch1-3 showed excitatory influence over the vmPFC, PCC, left IPL and VP and an inhibitory influence on the bilateral anterior insula and dACC. Conversely, Ch4 illustrated an inhibitory influence over the vmPFC, PCC, bilateral IPL and dACC, and an excitatory influence over the bilateral anterior insula and VP. VP showed a similar connectivity pattern to Ch4, with an excitatory influence over the bilateral anterior insula and Ch4, and an inhibitory influence over the vmPFC, PCC, and bilateral IPL.

### Effective Connectivity of Differences in Clinical Participants Compared with Healthy Controls

Clinical participants demonstrated strong evidence for alterations in effective connectivity, particularly greater inhibitory connectivity originating from Ch4 to regions of the DMN and from VP to posterior DMN regions (*Fig. 2B*).

Clinical participants were shown to have greater inhibitory connectivity from Ch4 to the vmPFC, dACC, PCC, and bilateral IPL (*Fig. 2C*). Greater excitatory connectivity from Ch4 to the bilateral anterior insula was also observed. Clinical participants illustrated greater inhibitory connectivity from the VP to the PCC, and bilateral IPL. Afferent connections to Ch4 also showed between group differences, specifically from the PCC to Ch4 and from Ch1-3 to Ch4. Notably, greater self-inhibition of vmPFC and reduced self-inhibition of Ch4 were identified in our clinical group.

To ensure that these effects were not confounded by the presence of other conditions in our transdiagnostic sample we conducted a subgroup analysis including only those participants with an anxiety or depressive disorder. These results demonstrated similar effects as the whole sample analysis, with consistent effects identified from Ch4 to the vmPFC, dACC, PCC, VP, the self-connection of the vmPFC and Ch4, as well as from Ch1-3 and the PCC to Ch4 (*Supplementary Fig. S2*).

### Parametric Empirical Bayes Cross-validation and Prediction of Clinical Diagnosis

Having identified multiple between group effects, we then explored whether the size of these effects was sufficiently large to be clinically useful using leave-one-out cross-validation (LOOCV) in the PEB framework. To restrict the number of comparisons made, we limited this to effects that were identified both in the analysis including all clinical participants and the subgroup analysis with only those with depressive and anxiety disorders. Of these connections, effective connectivity from Ch4 to the vmPFC (*r* = 0.22, *p* = 0.003), dACC (*r* = 0.14, *p* = 0.049), VP (*r* = 0.24, *p* = 0.002) and from the PCC to Ch4 (*r* = 0.14, *p* = 0.042) were all observed to significantly predict group. Using the altered connectivity of these connections together resulted in an overall correlation of *r* = 0.32, *p* < 0.001 (*Fig. 3*). This is consistent with an area under the curve of 0.67 (95% CI: 0.58-0.75), with a sensitivity of 0.57 and specificity 0.72 (*Supplementary Fig. S3*). Thus, the difference across the groups in effective connectivity was sufficiently large to predict whether a left-out subject was a healthy control or clinical participant above chance, however, much of the variability was left unexplained.

**Fig. 3.**
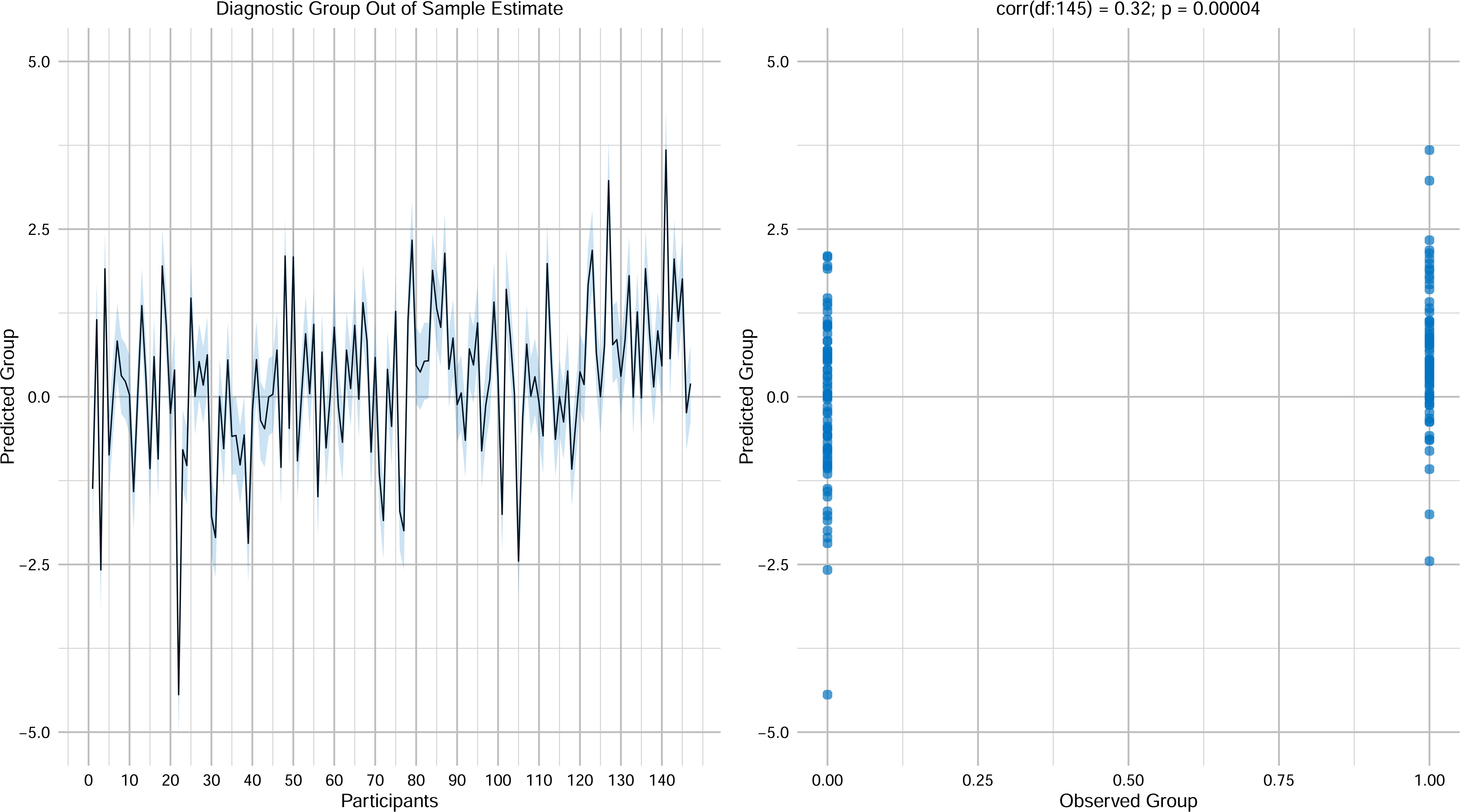
Leave-one-out cross-validation predicting group category for each participant. **Left:** The out-of-samples estimate of the group category (center line) with 90% credible interval (shaded area). **Right:** The Pearson’s correlation (one-tailed) between observed scores and the expected values for each individual participant using altered connections. Healthy controls are shown on the left (x = 0) and clinical participants on the right (x = 1).

### Relationship between Effective Connectivity and Depressive and Anxiety Symptoms in Clinical Participants

Following our identification of these between group differences, we examined whether these parameters similarly related to depressive and anxiety symptom severity for those with an anxiety or depressive disorder (*Fig. 4*). Notably, the connectivity alterations related to symptom severity largely did not overlap with those associated with clinical diagnosis.

**Fig. 4.**
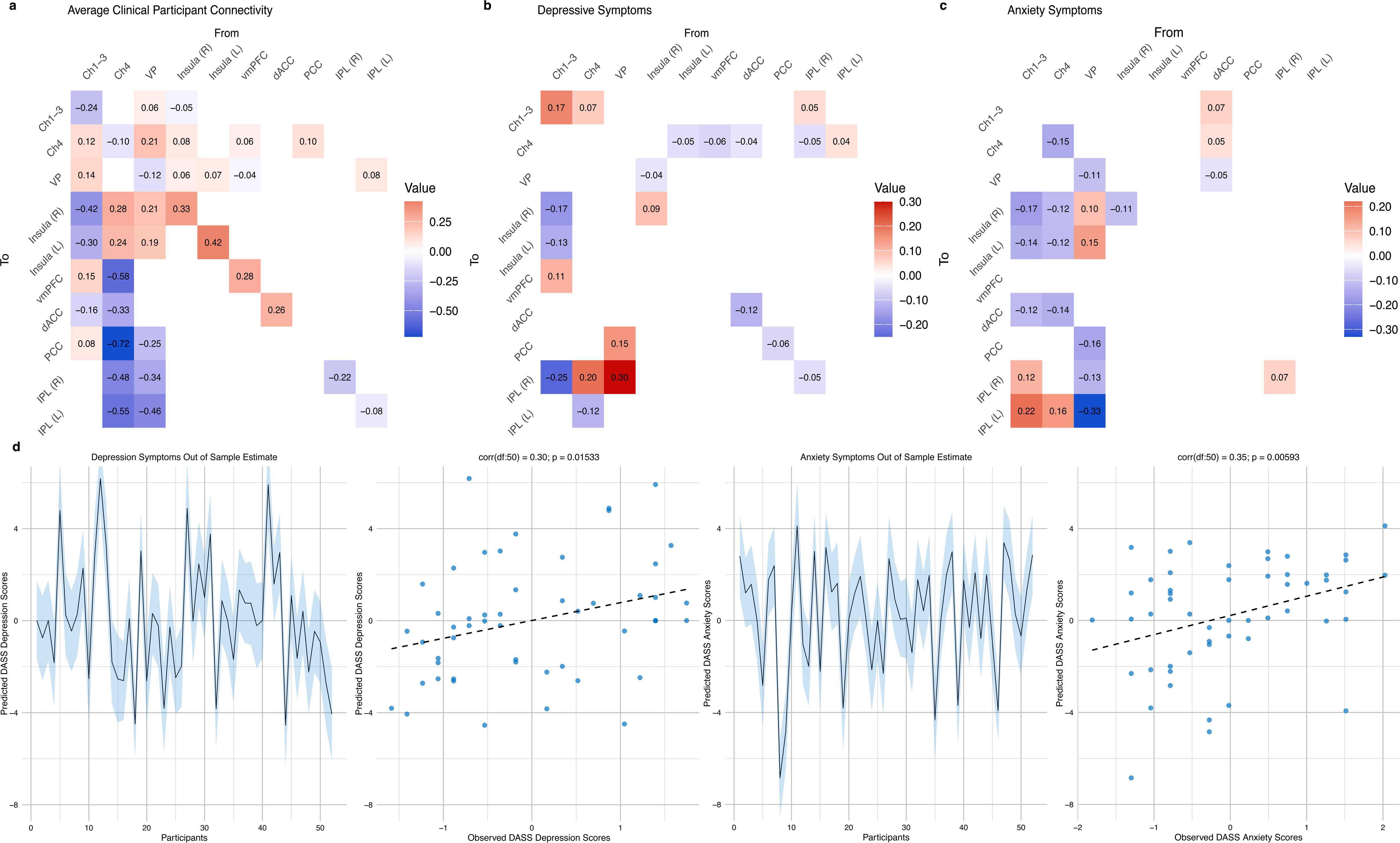
Association between effective connectivity and depressive and anxiety symptom severity. Adjacency matrices show **a**, the average effect across participants with depressive and anxiety disorder diagnoses (*n* = 52), **b**, changes associated with depressive symptom severity, **c**, changes associated with anxiety symptom severity. For **a**, red cells indicate excitatory connections, whereas blue cells indicate inhibition. For **b**, and **c**, these values add (red) to or subtract (blue) from the connectivity in **a**,. Leave-one-out cross validation results indicate that connectivity from **d**, Ch1 to the right insula, from VP to the right IPL, and the dACC self-inhibition was predictive of depressive symptom severity, whereas connectivity from **e**, Ch1-3 to right IPL and insula was predictive of anxiety symptom severity. Left (**d**, and **e**,): The out-of-samples estimated of the z-score DASS severity (center line) with 90% credible interval (shaded area). Right (**d**, and **e**,): The Pearson’s correlation (one-tailed) between observed scores and the expected values for each individual participant using altered connections. *Ch1-3* medial septal nucleus, *Ch4* nucleus basalis of Meynert*, dACC* dorsal anterior cingulate cortex, *DASS* Depression Anxiety Stress Scale*, IPL* inferior partial lobule, *PCC* posterior cingulate cortex, *vmPFC* ventromedial prefrontal cortex, VP ventral pallidum.

#### Depressive symptom severity associations

Depressive symptom severity was associated with alterations largely stemming from Ch1 and to Ch4 (*Fig. 4B*). Specifically, greater depressive symptom severity was observed to be associated with greater inhibition from Ch1-3 to the bilateral anterior insula, greater excitation from Ch1-3 to the vmPFC, and reduced excitation from Ch1-3 to the right IPL. Greater excitation to Ch1-3 was identified from Ch4 and from the right IPL.

Reduced inhibition was observed from Ch4 to the right IPL and greater inhibition from Ch4 to the left IPL. Reduced excitation to Ch4 was observed from the left insula, vmPFC, dACC, and left IPL, whereas greater excitation was observed from the left IPL. Reduced inhibition was also illustrated from VP to the PCC and IPL, as well as to the VP from the right anterior insula. Reduced self-inhibition of the dACC, PCC, and right IPL and greater self-inhibition of Ch1-3 and right insula were also observed.

We then tested whether parameters greater than 0.10 were able to predict symptom severity using PEB LOOCV. For depressive symptoms, only connectivity from Ch1-3 to the right insula (*r* = 0.29, *p* = 0.018), from VP to the right IPL (*r* = 0.24, *p* = 0.044), and the dACC self-inhibition (*r* = 0.27, p = 0.027) were significantly predictive, for an overall correlation of *r* = 0.30, *p* = 0.015 (*Fig. 4D*).

#### Anxiety symptom severity associations

Anxiety symptom severity was conversely largely associated with broad alterations across the three basal forebrain regions, particularly in their relationships with the anterior insula and dACC (*Fig. 4C*). Anxiety symptom severity was associated with greater inhibition from Ch1-3 to the bilateral anterior insula, and dACC, and greater excitation from Ch1-3 to the bilateral IPL. Conversely, reduced excitation from Ch4 to the bilateral anterior insula, greater inhibition from Ch4 to the dACC, and reduced inhibition from Ch4 to the left IPL was noted. Greater excitation from VP to the bilateral anterior insula and greater inhibition to the PCC and bilateral IPL were also observed. Top-down alterations from the dACC were also observed, specifically greater inhibition of the VP and greater excitation of Ch1-3 and Ch4. Reduced self-inhibition was observed of Ch4, VP, and right insula, whereas greater self-inhibition was shown in the right IPL.

As above, we then tested whether the parameters were able to predict symptom severity using PEB LOOCV. Connectivity from Ch1-3 to left IPL (r = 0.30, p = 0.014) and insula (r = 0.27, p = 0.026) as well as from VP to left IPL (r = 0.25, P = 0.006) were significantly predictive of anxiety symptom severity, for an overall correlation of *r* = 0.35, *p* = 0.006 (*Fig. 4E*).

## Discussion

This study examined alterations in basal forebrain resting-state effective connectivity across depressive and anxiety disorders. Our findings highlight the differential roles of Ch1-3, Ch4, and VP in coordinating human intrinsic brain networks. We identified that people with depressive and anxiety disorders demonstrated altered regulation of the DMN and salience networks from basal forebrain subregions, particularly Ch4. Our results therefore both support the basal forebrain’s role in the coordination of the DMN and salience network and the broad dysfunction of this circuitry in depressive and anxiety disorder patients.

The basal forebrain contains a heterogeneous population of neurons, including GABAergic, cholinergic, glutamatergic, and peptidergic neurons^42–44^, which interact locally in addition to projecting cortically^45^. Parvalbumin positive GABAergic neurons have emerged as likely candidates for the anatomical substrate of the basal forebrain responsible for DMN regulation^46,47^. These GABAergic neurons have been shown to project to major DMN nodes, including the anterior cingulate and retrosplenial cortices^48,49^. Optogenetic work has specifically identified causal evidence for the critical role of parvalbumin positive neurons in triggering state switching by activating the DMN and inducing associated behaviors^48^. These effects are also consistent with observations from VP GABAergic neurons, with stimulation of these neurons being shown to automatize internally focused behavior and impair attention directed towards external stimuli^25^. This function mirrors increased ruminative thought observed across depressive and anxiety disorders^50^. Moreover, local optogenetic inhibition of VP GABAergic neurons triggers transitions from quiet wakefulness to exploratory behavior, with associated cortical activation being partially dependent on cholinergic transmission^51^. Acetylcholine release and activity from cholinergic neurons in the basal forebrain have also been shown to be associated with reductions in DMN activity and connectivity^18,52^. Together this indicates that regulation of the DMN likely occurs through both cholinergic and non-cholinergic mechanisms within the basal forebrain, consistent with findings from our current study.

Previous research has reported greater negative connectivity from the anterior DMN to regions of the salience and central executive networks in MDD^9^, indicating an increased influence of the DMN over these networks^53^. These effects are congruent with research identifying that MDD is characterized by greater time spent in a coactivation pattern involving both activation of the DMN and deactivation of the frontoparietal network^54^. This DMN dominance has been hypothesized to underlie a dysfunctional ability to shift attention from self-directed thoughts to external, goal-related behavior in MDD^55,56^. Notably, the results of the current study and others indicate that these effects are directly influenced by changes in basal forebrain activity^14,16,25^. The fact that the largest effects were observed in regards to Ch4 connectivity is notable given its high proportion of cholinergic neurons^19^. The cholinergic basal forebrain system has been hypothesized to modulate information processing and attention by enhancing functional segregation within the brain^57^. This is further corroborated by recent research which has shown that local inactivation of Ch4 leads to a decrease in the energy barriers required for a brain state transition in cortical activity, thereby increasing the likelihood of transitioning between different brain states^58,59^. Dysfunction of the cholinergic system has been implicated in the development of cognitive impairments consistent with depressive and anxiety symptoms, including attentional deficits, memory dysfunction, and negativity biases^60,61^. As such, a combination of abnormal influences from both the VP and Ch4 may result in altered DMN activity and an impaired ability to switch between network states^14^, which in turn contributes to the development of depressive and anxiety symptoms.

Dysfunctional Ch4 connectivity across these disorders may have direct implications for treatment using medications that affect cholinergic transmission. Scopolamine is a nonspecific muscarinic receptor antagonist which has demonstrated some evidence as a rapid antidepressant and anxiolytic, with effects occurring as quickly as 3 days post-infusion^62–64^. However, there have been relatively few high-quality trials investigating scopolamine’s antidepressant effects and these studies have demonstrated inconsistent evidence for its effectiveness due to differences in administration routes and dosage^63,65^. An additional contributor to this response variability may include dysfunction in the Ch4 circuitry observed within the current study, and as such, be indicative of a specific subtype with sensitivity to scopolamine treatment. Relatively few investigations have been undertaken to identify potential predictors of scopolamine treatment response^66^, with fewer still investigating neuroimaging predictors. However, growing preclinical evidence indicates that scopolamine’s rapid antidepressant effects, akin to other rapid antidepressant such as ketamine, occurs via inhibition of GABAergic interneurons in the mPFC^67^. As dysfunction both within and between Ch4 and vmPFC were identified above, prospective investigation into the utility of these connectivity alterations in predicting response to scopolamine is warranted. An improved understanding of the neurobiological mechanisms involved in scopolamine treatment and its effects on basal forebrain circuitry may provide clinical guidance for future applications of this treatment.

## Limitations

While the transdiagnostic nature of the clinical group under investigation is a strength of the study and is representative of community samples, the non-specific nature of the sample impairs the specificity and potential clinical utility of the work. Using discrete diagnostic categories to investigate this circuitry and its relationship with symptoms may be more useful in identifying specific clinical targets. While 7T fMRI has many advantages over 3T, it also has several notable drawbacks including increased susceptibility artifacts which may have contributed to signal dropout in the basal forebrain resulting in the exclusion of several participants^68^. Given the role of the cholinergic system in modulating arousal and attention^69^, physiological measure such as eye tracking may have provided greater insight into the discrete time periods in which this circuitry was most active and as such should be integrated into future research. It is unclear whether the changes highlighted here represent a trait or state marker of these disorders, however, longitudinal research examining the stability of these parameters as well as the presence of these changes in those at risk for depression and anxiety may aid in clarifying these effects. Whilst beyond the scope of this study, it should be noted that the amygdala and basal forebrain demonstrate dense reciprocal connections with one another^70,71^. Given that this pathway has been implicated in chronic stress models^72^ and depression-like behavior in rodents^73^ it may be worth examining in future studies.

## Conclusions

Within this study we have presented evidence for effective connectivity alterations across basal forebrain subregions in depressive and anxiety disorder patients. This work supports the notion that the basal forebrain is an important functional regulator of intrinsic brain networks and that different basal forebrain subregions have specific influences over these brain areas. We identified widespread increased inhibition from Ch4 to regions of the DMN and the dACC. Similar alterations were observed from the VP to regions of the posterior DMN. While a putative cholinergic basis of the aforementioned dysfunction needs to be established through future research, this may highlight novel treatment targets for depressive and anxiety disorders. Future work directly examining the effect of cholinergic receptor antagonists on basal forebrain circuitry in parallel with their effect on depressive and anxiety symptoms would aid in elucidating this relationship.

## Methods

A transdiagnostic sample of 79 clinical participants were recruited to the study from The University of Melbourne Psychology Clinic and via the Research Experience Program at the Melbourne School of Psychological Sciences. All clinical participants underwent the Diagnostic Interview for Anxiety, Mood, and OCD and Related Neuropsychiatric Disorders (DIAMOND)^74^, a semi-structured clinical interview based on DSM-5 criteria, to identify those who met criteria for a mood and/or anxiety disorder. Participants were also included if they met criteria for stress and/or eating disorders due to the high level of depressive and anxiety symptoms present in these conditions^75,76^. Participants who met criteria for psychosis, bipolar disorder, paraphilic disorder, dissociative disorder, or conduct disorder were excluded. Clinical participants were compared to 92 healthy control participants recruited via publicly accessible online advertisements. These participants had no current or past diagnosis of a mental health disorder as assessed through the Mini-International Neuropsychiatric Interview-7^77^. All participants were between 18 to 40 years of age, fluent English speakers, had no MRI contraindications including pregnancy, and had normal or corrected-to-normal visual acuity. All participants provided written informed consent prior to the study, following a complete description of the study protocol. This study and its protocol were approved by the Melbourne Health Human Research and Ethics Committee. All participants attended a single 7T scanning session at the Melbourne Brain Imaging Unit (The University of Melbourne, Parkville), were recruited between October 2021 and December 2023, and were adequately reimbursed for their time. The DASS was administered prior to the session to provide an overall measure of negative mood states, as well as sub-scales scores of depression, anxiety, and stress-tension^78^. Of the original sample, three participants were excluded due to data quality issues (three clinical participants), five participants were excluded due to excessive head movement during scanning (four controls and one clinical participant), and 16 due to insufficient activity mapping across all regions necessary for DCM (11 controls and five clinical participants; for further details see Times Series Extraction and Model Specification below). This resulted in 70 clinical participants and 77 healthy controls being included in the primary analysis.

### Image Acquisition

MRI acquisition was performed on a Siemens 7T Plus research scanner (Siemens Healthcare, Erlangen, Germany) equipped with a 32Rx/1Tx channel head coil (Nova Medical Inc., Wilmington MA, USA). Functional (T2*-weighted) images were obtained using a multi-band and grappa accelerated gradient-echo planar imaging sequence^79^ in the steady state (multi-band factor, 6; parallel acceleration factor, 2; repetition time, 800 ms; echo time, 22.2 ms; flip angle, 45°) in a 20.8 cm field-of-view, with a 130×130-pixel matrix, a slice thickness of 1.6mm (no gap) and in-plane voxel size=1.6×1.6 mm. Eighty-four interleaved slices were acquired parallel to the anterior-posterior commissure line, covering the whole brain. The resting-state sequence duration was 5 minutes and 20 seconds, corresponding to 400 whole-brain echo-planar imaging volumes. Participants were instructed to keep their eyes closed but not to fall asleep for the duration of the scan.

A high resolution structural T1-weighted image was obtained from each participant using magnetization-prepared 2 rapid gradient echo sequence (MP2RAGE^80^) for co-registration with the functional images (parallel reduction factor, 4; repetition time, 5 seconds; echo time, 2.04 ms; flip angle, 13°) in a 24 cm field of view, with a 330×330–pixel matrix, in-plane voxel size = 0.75×0.75 mm, slice thickness = 0.75 mm with 224 sagittal slices aligned parallel to the midline. To minimize head movement during scanning, foam pads were inserted to the either side of participants’ heads. Respiration and cardiac pulse were also recorded during the session at 50 Hz and 200 Hz, respectively, using a respiratory belt and pulse-oximeter (Siemens, Germany) to be used for physiological noise correction.

### Preprocessing

Imaging data were preprocessed using Statistical Parametric Mapping (SPM) 12 (v7771, Welcome Trust Centre for Neuroimaging, London, UK) within a MATLAB 2023b environment (The MathWorks Inc., Natick, MA) on the Spartan High Performance Computer hosted at The University of Melbourne^81^. Motion artifacts were corrected by realigning each participant’s timeseries to the mean image, and all images were resampled using 4th Degree B-Spline interpolation. Participants were excluded if movement exceeded a mean total displacement of 1.6mm (∼1 native voxel) as assessed through the Motion Fingerprint toolbox^82^.

Each participant’s anatomical images (denoised MP2RAGE^83^) were co-registered to their respective mean functional image, segmented, and normalized to the International Consortium of Brain Mapping template using the unified segmentation and the Diffeomorphic Anatomical Registration Through Exponentiated Lie Algebra approach^84^. Smoothing was applied with a 3.2-mm^3^ full-width-at-half-maximum Gaussian kernel to preserve spatial specificity. Physiological noise was modeled using PhysIO toolbox^85^. Cardiac and respiratory recordings were imported to the toolbox, in which physiological noise models were applied to these recordings (i.e., the retrospective image-based correction for a third order cardiac, fourth order respiratory, and first order interaction Fourier expansion of cardiac and respiratory phase^86^). Normalized white matter and cerebrospinal fluid tissue segmentations from each participant’s structural image were used to construct noise regions of interest from which mean timeseries and principal components were extracted as nuisance regressors using CompCor^87^. These regressors were added to six movement regressors estimated through realignment and taken to the first-level general linear model to remove as sources of variance that were not of interest.

### Timeseries Extraction and Model Specification

Volumes-of-interest (VOIs) from each of our regions of interest were defined in SPM12 by calculating the principal eigenvariate of all voxels within a given part of the brain. To extract the basal forebrain subregions, we used basal forebrain masks based on the stereotaxic probabilistic maps of the magnocellular basal forebrain space by Zaborszky and colleagues^88^ for Ch1-3 and Ch4, and the subcortical mask from Pauli and colleagues^89^ for the VP (*Fig. 1*). Seed-to-whole brain functional connectivity analyses were conducted to ensure that these masks accurately captured the intended regions and their associated functional connectivity patterns (Supplementary Methods and Results). The Montreal Neurological Institute coordinates for core regions of the DMN and salience network were identified using previously reported peak coordinates from meta-analyses and large-scale functional connectivity studies. The precise coordinates were as follows: vmPFC [-3, 39, -2]^90^, PCC [-2 -36 37]^90^, IPL [44 -65 32; -40 -66 32]^91^, anterior insula [36, 16, 4; -35, 14, 5]^92^, dACC [4, 30, 30]^93^. The timeseries from these VOIs were extracted using the principal eigenvariate of all voxels in a sphere within a radius of 6 mm of the centroids for midline regions (vmPFC, PCC, dACC) and 4 mm for clearly lateralized regions (IPL and anterior insula; *Fig. 1*). To avoid potential oversampling of midline voxels, single VOIs were used for these regions, consistent with past DCM studies^9,94,95^.

While traditional deterministic and stochastic DCM operates by analyzing the timeseries data directly, spectral DCM estimates effective connectivity using the cross-spectral density (for further details see Supplementary Methods). This is a second-order summary statistic of the timeseries data and thus allows the estimation of effective connectivity parameters for resting-state data with a greatly increased computational efficiency^96^. Our model space was specified using DCM 12.5 with default priors. To address our hypotheses concerning the distinct effects of basal forebrain subregions, we modeled bidirectional connectivity to and from these subregions to all cortical regions highlighted above.

### Parametric Empirical Bayes

To examine the impact of group differences on individual parameters, we employed PEB^97^. Unlike comparisons using classical frequentist tests which only use estimated mean values, PEB allows for the inclusion of estimated variance (in addition to the estimated means) of each parameter when investigating between-group effects. The PP was calculated using the free energy (with vs. without) method. This approach compares the evidence for all models in which a particular parameter is switched on with the evidence for models in which it is switched off. A PP threshold of greater than 0.99 was used to identify those parameters which had a *very strong* amount of evidence^39^. We constructed a second level PEB model including four regressors. The first regressor modeled the average connectivity observed across healthy control participants, with subsequent regressors either increasing or decreasing this baseline connectivity. The subsequent regressors included the difference between clinical participants and healthy controls, as well as age and gender as covariates. The same process was repeated for the sub-analysis examining symptom severity in the clinical group, however, in this case the first regressor represented the average connectivity across the clinical group, with subsequent regressors representing depressive symptom severity, anxiety symptom severity, age, and gender. Having estimated a group level PEB model, we then searched over nested PEB models using Bayesian model reduction pruning parameters that did not contribute to overall model evidence^98,99^. Bayesian model averaging was performed on these models after the final iteration to determine the strength of connections in the last Occam’s window of 256 models.

### Leave-one-out Cross-validation

To assess the predictive validity of parameters which demonstrated between-group differences, we used LOOCV across all participants. In its implementation within the PEB framework (see^39^ for more details), LOOCV aims to determine whether the size of between-group effects on parameters is sufficiently large to predict a variable of interest^39^. In this case, this was employed to determine whether identified parameters could predict whether an individual was a healthy control or clinical participant. A group-level PEB model was estimated for all participants while excluding one participant, then this PEB model was used to predict the left-out subject’s group. Predicted group membership was then correlated with the observed group. A significant correlation between the expected and observed values demonstrates that the effect size was sufficiently large to predict the left-out subjects’ group above chance. We also repeated this procedure to investigate depression and anxiety symptom severity in the clinical group.

### Additional Statistics

Analyses of between-group differences for clinical and demographic characteristics were calculated in SPSS version 30 (IBMCorp., Armonk, NY), using two-tailed significance tests. Comparisons were adjusted for multiple comparisons using the Benjamini–Hochberg correction^100^ to determine significance (*p* < 0.05). No statistical method was used to predetermine sample size.

## Data availability

Individual participant and group level effective connectivity data for this study are available at: https://github.com/alecJamieson/basal_forebrain_EC/tree/main/Data.

## Code availability

MATLAB and R scripts used to generate the 2^nd^ level results and associated figures of this study are also available at: https://github.com/alecJamieson/basal_forebrain_EC.

## Acknowledgements

The authors thank Braden Thai, Holly Carey and Amy Nielson for their contributions to data collection. The authors also acknowledge the facilities and scientific and technical assistance of the National Imaging Facility, a National Collaborative Research Infrastructure Strategy (NCRIS) capability, at the Melbourne Brain Centre Imaging Unit, University of Melbourne. In addition, the authors are grateful to Siemens for providing the MP2RAGE sequence as a “works in progress package” and CMRR (University of Minnesota) for sharing the multiband EPI sequence.

## Author Contribution Statement

AJJ: Conceptualization, Formal analysis, Methodology, Writing - original draft, Visualization. TS: Conceptualization, Funding acquisition, Software, Writing - review & editing. CGD: Writing - review & editing. SI: Investigation, Methodology, Writing - review & editing. JAA: Investigation, Writing - review & editing. BAM: Funding acquisition, Writing - review & editing. RKG: Investigation; Writing - review & editing. KLF: Resources, Writing - review & editing. BJH: Conceptualization, Funding acquisition, Methodology, Supervision.

