## Supplementary Materials for "Altered basal forebrain regulation of intrinsic brain networks in depressive and anxiety disorders"

**Supplementary Methods……………………………………………………………………2**

**Supplementary Results………………………………………..…………………………5**

**Supplementary Tables………………………………………………………………………6**

**Supplementary Figures……………………………………………………………………..11**

### Supplementary Methods

#### Functional Connectivity of Basal Forebrain Regions

To confirm that the basal forebrain masks were capturing the intended regions and their associated functional connectivity patterns, we conducted a seed-to-whole brain functional connectivity analyses in the healthy controls using the CONN Toolbox[1]. Using conventional conservative thresholds (whole-brain FWE corrected, $p < .001$), connectivity patterns were observed across the cortex. To isolate the regions which demonstrated the most robust connectivity effects, we applied a further threshold that restricted results to the top 5% of voxels. This was equivalent to applying t-thresholds of 11.49 for the VP, 10.58 for Ch4, and 8.48 for Ch1-3.

#### Spectral Dynamic Causal Modelling

Dynamic causal modelling (DCM) is Bayesian framework that infers the directed (causal) connectivity among the neuronal systems – referred to as effective connectivity. We recently proposed a new DCM for resting state fMRI – based upon a deterministic model that generates predicted cross spectra – referred to as spectral DCM. In order to model resting state activity – in the absence of external stimuli – we will have to add a stochastic component, i.e. neural fluctuations, to the classical DCM based on ordinary differential equations. Mathematically, we can express the formulation of the stochastic generative model using a set of two equations. First is the neuronal state equation, namely:

$$\dot{x}(t) = f(x(t), u(t), \theta) + v(t) \tag{1}$$

and second is the observation equation, which is a static nonlinear mapping from the hidden physiological states in (1) to the observed BOLD activity and is written as:

$$y(t) = h(x(t), \varphi) + e(t) \tag{2}$$

where $\dot{x}(\mathrm{t})$ is the rate of change of the neuronal states $x(t)$, $\theta$ are unknown parameters (i.e. the effective connectivity) and $v(t)$ (resp. $e(t)$) is the stochastic process – called the state noise (resp. the measurement or observation noise) – modelling the random neuronal fluctuations that drive the resting state activity. In the observation equations, $\varphi$ are the unknown parameters of the (haemodynamic) observation function and $u(t)$ represents any exogenous (or experimental) inputs that drive the hidden states – that are usually absent in resting state designs[2]. Spectral DCM furnishes a constrained inversion of the stochastic model by parameterizing the neuronal fluctuations $v(t)$. Spectral DCM simplifies the generative model by replacing the original timeseries with their second-order statistics (i.e., cross spectra). This means, instead of estimating time varying hidden states, we are estimating their covariance which is time invariant. Then we simply need to estimate the covariance of the random fluctuations; where a scale free (power law) form for the state noise (resp. observation noise) is used – motivated from previous work on neuronal activity [3-5] – as follows:

$$g_v(\omega, \theta) = \alpha_v \omega^{-\beta_v} \qquad (3)$$

Here, $\{\alpha, \beta\} \subset \theta$ are the parameters controlling the amplitudes and exponents of the spectral density of the neural fluctuations. The parameterisation of endogenous fluctuations means that the states are no longer probabilistic; hence the inversion scheme is significantly simpler, requiring estimation of only the parameters (and hyperparameters) of the model. We used standard Bayesian model inversion to infer the parameters of the model in (1), (2) and (3), from the observed signal $y(t)$. The description of the Bayesian model inversion procedures based on variational Laplace can be found elsewhere for the interested readers [6-8]. Scatterplots and correlations between the expected values of connectivity parameters originating from basal forebrain subregions and depressive and anxiety symptom severity are depicted *Supplementary Fig. S4 and S5.*

**Parametric Empirical Bayes**

Empirical Bayes refers to the Bayesian inversion or fitting of hierarchical models. In hierarchical models, constraints on the posterior density over model parameters at any given level are provided by the level above. These constraints are called empirical priors because they are informed by empirical data. A second-level or between-subjects model over parameters has recently been introduced, which represents how individual (within-subject) connections derive from the subjects' group membership[9] – based on parametric empirical

Bayes (PEB). This approach calls on Bayesian Model Reduction (BMR) to finesse the inversion of multiple models of a single dataset or a single (hierarchical) model of multiple datasets. BMR allows one to compute posterior densities over model parameters, under new prior densities, without explicitly inverting the model again. For example, one can invert a DCM for each subject in a group and then evaluate the posterior density over group effects, using the posterior densities over parameters from the single subject inversion. This may improve subject-specific parameter estimates, by using group-level estimates to rescue individual DCM from local optima. Mathematically, for DCM studies with *N* subjects and *M* parameters per DCM, we have a hierarchical model, where the responses of the $i$-th subject and the distribution of the parameters over subjects can be modeled as:

$$y_i = \Gamma_i^{(1)}(\theta^{(1)}) + \varepsilon_i^{(1)} \quad (4)$$

$$\theta^{(1)} = \Gamma^{(2)}\big(\theta^{(2)}\big) + \varepsilon^{(2)}$$

$$\theta^{(2)} = \eta + \varepsilon^{(3)}$$

where, $y_i$ is the BOLD time series from *i-th* subject *and* $\Gamma_i^{(1)}$ is a nonlinear mapping from the parameters of a model to the predicted response $y$ for e.g. as shown in Eq. S1 above. $\varepsilon_i^{(1)}$ is independent and identically distributed (i.i.d.) observation noise (equivalent to $e(t)$ in Eq. S2). In this hierarchical form, *empirical priors* encoding second (between-subject) level effects place constraints on subject-specific parameters. The second level would be a linear model where the random effects are parameterized in terms of their precision:

$$\Gamma^{(2)}\big(\theta^{(2)}\big) = (X \otimes W)\beta \quad (5)$$

where, $\beta \subset \theta$ are group means or effects encoded by a design matrix with between $X$ and within-subject $W$ parts. The between-subject part encodes differences among subjects or covariates such as age, while the within-subject part specifies mixtures of parameters that show random effects. We assume that the first column of the design matrix is a constant term, modelling group means and subsequent columns encode group differences or covariates such as age.

**Magnitude of Effect Size**

Currently there is no standard heuristics for determining the size of effects present in DCM studies. To aid in the interpretation of our findings and future studies we have devised a set of standard ranges based on magnitude of the posterior expectations (means) in relation to the priors of the model. Greater effects are necessary to move posterior expectations further away from the prior mean of zero. Given the default priors with a mean of zero and a variance of 1/64, we would expect that 90% of the values would be ±0.21 of the mean, 99% of the values would be ±0.32 of the mean, 99.99% of values fall within ±0.49 of the mean. As such, these ranges can be used to assess the size of the effects demonstrated here and in other studies (Supplementary Table S1). Notably, these ranges are specifically calculated based on the prior distribution present for intrinsic connections and as such different ranges would need to be calculated for modulatory and self-connections.

**Supplementary Results**

**Functional Connectivity of Basal Forebrain Subregions**

Seed-to-whole brain functional connectivity analyses in healthy controls revealed results consistent with effects observed in previous functional connectivity investigations of basal forebrain connectivity [10,11]. Ch1-3 demonstrated significant functional connectivity with regions of the default mode network including the ventromedial prefrontal cortex, posterior cingulate cortex extending to the retrosplenial cortex and hippocampus, bilateral inferior parietal lobules, and frontal eye fields (*Supplementary Fig. S1A*). Conversely, Ch4 demonstrated functional connectivity with regions including the dorsal anterior and posterior insula, pregenual extending to the dorsal anterior (dACC) and mid-cingulate cortex, dorsolateral prefrontal cortex, primary motor cortex, supplementary motor area, thalamus, and cerebellum (*Supplementary Fig. S1B*). Similar to Ch4, the VP demonstrated functional connectivity across cortical midline structures including pregenual ACC, dACC, mid-cingulate cortex, and PCC extending to the precuneus, as well as primary motor cortex, supplementary motor area, fusiform gyrus, cerebellum, and thalamus (*Supplementary Fig. S1C*). The full list of functional connectivity results is given in Supplementary Tables S2 (Ch1-3), S3 (Ch4), and S4 (VP).

Supplementary Table S1

*Effect Size Magnitude Scale for Intrinsic between Region DCM Parameters*

| Size | Parameter Value |
|---|---|
| Small | < 0.21, > -0.21 |
| Medium | ±0.21-0.32 |
| Large | ±0.32-0.49 |
| Very Large | > +0.49, < -0.49 |

Supplementary Table S2

*Resting-state Functional Connectivity of Ch1-3 in Healthy Controls*

| Brain region | BA | Coordinates | | | Cluster size ($1.6mm^3$ voxels) | *t*-value |
|---|---|---|---|---|---|---|
| | | X | Y | Z | | |
| Ch1-3 | - | 0 | 6 | -6 | 4189 | 28.35 |
| Rostral anterior cingulate | 24 | 4 | 30 | 0 | | 15.79 |
| Subgenual anterior cingulate | 25 | -2 | 24 | -2 | | 14.94 |
| Dorsal posterior cingulate | 31 | 4 | -62 | 24 | 4777 | 14.62 |
| Ventral posterior cingulate | 23 | -2 | -58 | 14 | | 14.3 |
| | | 2 | -40 | 32 | | 14.13 |
| Frontal eye fields | 8 | -22 | 38 | 42 | 455 | 11.95 |
| | | -20 | 28 | 44 | | 10.97 |
| | | -20 | 36 | 52 | | 10.77 |
| Inferior parietal lobule | 39 | 46 | -52 | 24 | 676 | 11.55 |
| | | 44 | -62 | 34 | | 10.94 |
| | | 34 | -76 | 36 | | 9.4 |
| Frontal eye fields | 8 | 24 | 36 | 46 | 544 | 11.49 |
| | | 20 | 28 | 38 | | 11.33 |
| | | 28 | 24 | 56 | | 9.76 |
| Inferior parietal lobule | 39 | -40 | -68 | 32 | 366 | 11.41 |
| | | -38 | -70 | 42 | | 11.25 |
| | | -44 | -62 | 40 | | 8.97 |
| Parahippocampal gyrus | 36 | -22 | -34 | -8 | 126 | 11.12 |
| Fusiform gyrus | 37 | -26 | -36 | -16 | | 10.18 |
| Cerebellum | - | -16 | -36 | -22 | | 9 |
| Midbrain | - | -14 | -20 | -12 | 48 | 11.04 |
| Parahippocampal gyrus | 36 | -20 | -18 | -18 | | 9.8 |
| Medial temporal gyrus | 21 | 60 | -4 | -16 | 140 | 10.71 |
| | | 56 | -14 | -14 | | 9.92 |
| Pars orbitalis | 47 | -22 | 26 | -12 | 58 | 10.69 |
| | | -28 | 30 | -16 | | 10.39 |
| Cerebellum | - | 22 | -84 | -24 | 554 | 10.57 |
| Fusiform gyrus | 37 | 46 | -56 | -22 | | 10.45 |
| | | 42 | -68 | -18 | | 10.1 |
| Cerebellum | - | 6 | -60 | -44 | 162 | 10.45 |
| | | -6 | -46 | -44 | | 10.07 |
| | | -2 | -66 | -36 | | 8.86 |
| Cerebellum | - | -22 | -80 | -28 | 116 | 10.22 |
| Visual association cortex | 19 | -32 | -80 | -20 | | 9.12 |
| Premotor cortex | 6 | 32 | -22 | 70 | 55 | 10.13 |
| Primary motor cortex | 4 | 26 | -30 | 76 | | 9.17 |

| | | | | | | |
|---|---|---|---|---|---|---|
| Supplementary motor cortex | 6 | -2 | 14 | 70 | 29 | 10.03 |
| Fusiform gyrus | 37 | -48 | -56 | -18 | 23 | 9.75 |
| Visual association cortex | 18 | -10 | -84 | -16 | 34 | 9.48 |
| | | -18 | -88 | -12 | | 8.59 |
| Primary motor cortex | 4 | 16 | -30 | 62 | 32 | 9.42 |
| Primary motor cortex | 4 | 4 | -30 | 80 | 20 | 9.4 |
| Supplementary motor cortex | 6 | 4 | -20 | 78 | | 9.17 |
| Primary sensory cortex | 1 | -4 | -34 | 80 | 21 | 9.36 |
| Primary visual cortex | 17 | 16 | -90 | 4 | 20 | 9.06 |

Supplementary Table S3

*Resting-state Functional Connectivity of Ch4 in Healthy Controls*

| Brain region | BA | Coordinates | | | Cluster size ($1.6mm^3$ voxels) | *t*-value |
|---|---|---|---|---|---|---|
| | | X | Y | Z | | |
| Ch4 | - | -20 | -2 | -12 | 4663 | 27.06 |
| | - | 18 | 0 | -12 | | 23.97 |
| Putamen | - | -20 | 6 | -4 | | 18.77 |
| Dorsal anterior cingulate | 32 | 6 | 14 | 34 | 2825 | 15.03 |
| Dorsal anterior cingulate | 24 | -4 | 16 | 26 | | 14.94 |
| | | -4 | 10 | 34 | | 13.98 |
| Cerebellum | - | 6 | -60 | -20 | 320 | 13.4 |
| | | -4 | -60 | -18 | | 12.28 |
| | | 6 | -58 | -28 | | 12.07 |
| Premotor cortex | 6 | 40 | -12 | 32 | 127 | 13.17 |
| Pars orbitalis | 47 | 28 | 30 | -12 | 71 | 12.99 |
| Dorsal anterior Insula | 13 | -38 | 8 | 0 | 139 | 12.62 |
| | | -34 | 24 | 2 | | 11.52 |
| Inferior frontal gyrus | 45 | -52 | 14 | -4 | 20 | 12.46 |
| Primary motor cortex | 4 | 20 | -32 | 62 | 168 | 12.37 |
| Primary sensory cortex | 1 | 22 | -32 | 76 | | 11.84 |
| Anterior prefrontal cortex | 10 | 2 | 52 | -8 | 30 | 11.8 |
| Orbitofrontal cortex | 11 | 4 | 32 | -8 | 30 | 11.7 |
| | | -8 | 34 | -10 | | 11.46 |
| Primary motor cortex | 4 | -18 | -32 | 66 | 78 | 11.53 |
| Primary sensory cortex | 1 | -24 | -30 | 74 | | 11.41 |
| Anterior prefrontal cortex | 10 | -32 | 46 | 34 | 49 | 11.48 |
| Dorsolateral prefrontal cortex | 17 | -24 | 36 | 26 | | 11.14 |
| Retrosplenial cortex | 30 | 0 | -42 | 20 | 27 | 11.19 |

Supplementary Table S4

*Resting-state Functional Connectivity of the Ventral Pallidum in Healthy Controls*

| Brain region | BA | Coordinates | | | Cluster size ($1.6mm^3$ voxels) | *t*-value |
|---|---|---|---|---|---|---|
| | | X | Y | Z | | |
| VP | - | 14 | 6 | -6 | 1398 | 26.8 |
| | | -14 | 6 | -6 | | 24.77 |
| Rostral anterior cingulate cortex | 32 | 8 | 28 | -4 | | 14.57 |
| Cerebellum | - | 6 | -60 | -26 | 1661 | 15.33 |
| | | -12 | -64 | -20 | | 14.64 |
| Fusiform gyrus | 37 | 44 | -64 | -18 | | 14.63 |
| Ventral posterior cingulate cortex | 23 | -10 | -60 | 10 | 686 | 14.14 |
| Cuneus | 7 | 26 | -78 | 36 | | 13.94 |
| Dorsal posterior cingulate cortex | 31 | 4 | -66 | 30 | | 13.92 |
| Dorsal anterior cingulate cortex | 32 | 2 | 6 | 34 | 1350 | 14.06 |
| Midcingulate cortex | 24 | 4 | -4 | 38 | | 13.62 |
| Ventral posterior cingulate cortex | 23 | 4 | -34 | 32 | | 13.53 |
| Fusiform gyrus | 37 | -46 | -56 | -20 | 96 | 14 |
| Visual association cortex | 19 | -42 | -74 | -14 | | 13.39 |
| Fusiform gyrus | 37 | -40 | -66 | -18 | | 12.13 |
| Primary auditory cortex | 41 | 52 | -18 | 8 | 53 | 13.63 |
| Parahippocampal gyrus | 36 | 28 | -16 | -24 | 46 | 13.56 |
| | | 26 | -24 | -18 | | 12.51 |
| Supplementary motor cortex | 6 | -2 | 6 | 74 | 52 | 12.9 |
| | | -4 | 6 | 66 | | 12.18 |
| | | 4 | 14 | 70 | | 11.98 |
| Visual association cortex | 18 | 6 | -88 | 20 | 43 | 12.88 |
| Primary visual cortex | 17 | 4 | -86 | 8 | | 12.16 |
| Precuneus | 7 | 26 | -58 | 52 | 67 | 12.88 |
| | | 28 | -66 | 56 | | 11.9 |
| Visual association cortex | 19 | -44 | -76 | -2 | 36 | 12.85 |
| Visual association cortex | 18 | -18 | -88 | 18 | 67 | 12.72 |
| Visual association cortex | 19 | -26 | -82 | 18 | | 12.27 |
| Dorsolateral prefrontal cortex | 9 | 26 | 42 | 34 | 23 | 12.66 |
| Visual association cortex | 18 | -18 | -94 | 8 | 20 | 12.66 |
| Primary motor cortex | 4 | 24 | -28 | 76 | 132 | 12.64 |
| Premotor cortex | 6 | 38 | -18 | 66 | | 12.28 |
| Primary sensory cortex | 1 | 20 | -36 | 66 | | 12.06 |
| Angular gyrus | 39 | -46 | -66 | 14 | 27 | 12.47 |

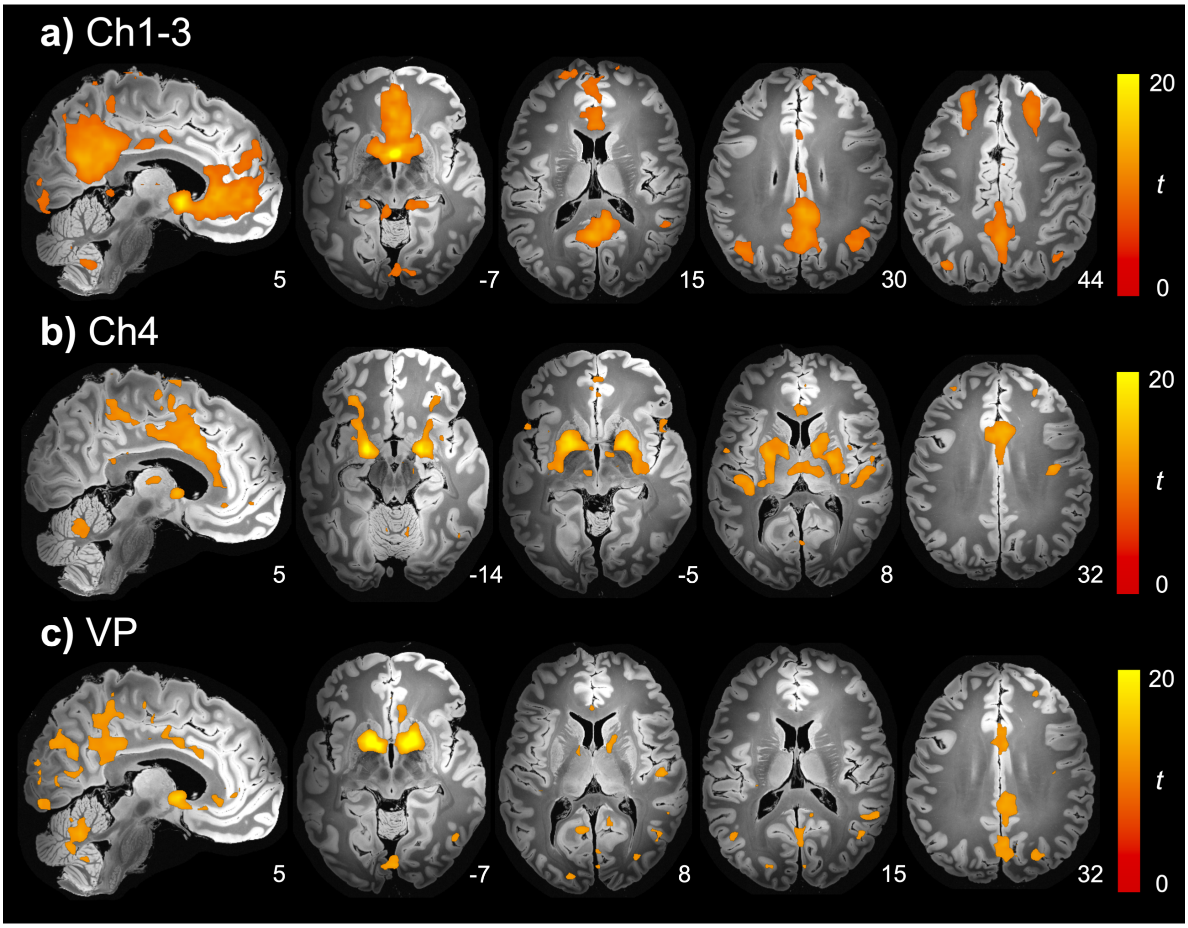


*Supplementary Fig. S1.* Functional connectivity of basal forebrain subregions in healthy controls. Functional connectivity **a**, of Ch1-3, **b**, Ch4, and **c**, ventral palladium. Results displayed for the top 5% of voxel t-values on the "Synthesized_FLASH25" (500 µm, MNI space) ex vivo template[12].

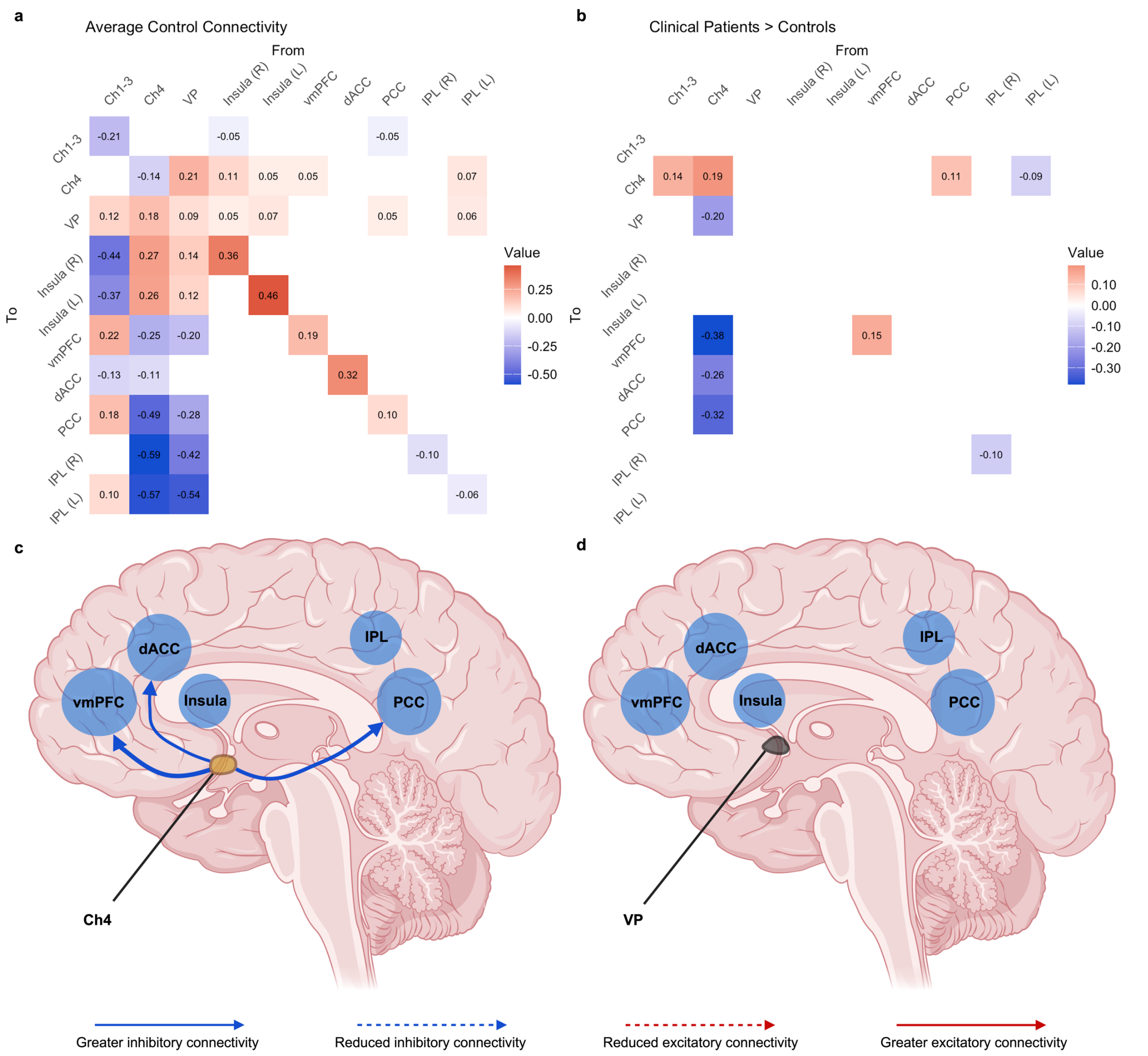


*Supplementary Fig. S2.* Effective connectivity differences identified in participants with depression and anxiety disorders. Adjacency matrices show the average effects observed in **a**, healthy controls (*n* = 77) and **b**, the difference between clinical participants (*n* = 52) and healthy controls. For **a**, red cells indicate excitatory connections whereas, blue cells indicate inhibitory connections. For **b**, these values add (red) to or subtract (blue) from the connectivity in **a**,. Cells representing connectivity between two regions are measured in hertz and diagonal cells indicate inhibitory self-connections which are unitless log scaling parameters. **c**, and **d**, illustrate differences in effective connectivity from the basal forebrain to cortical brain regions in clinical participants compared with healthy controls. Results indicate effects involving **c**, Ch4 (yellow) and **d**, ventral pallidum (black). Arrows have been weighted to indicate the relative size of the effects. Image created with BioRender (www.biorender.com).

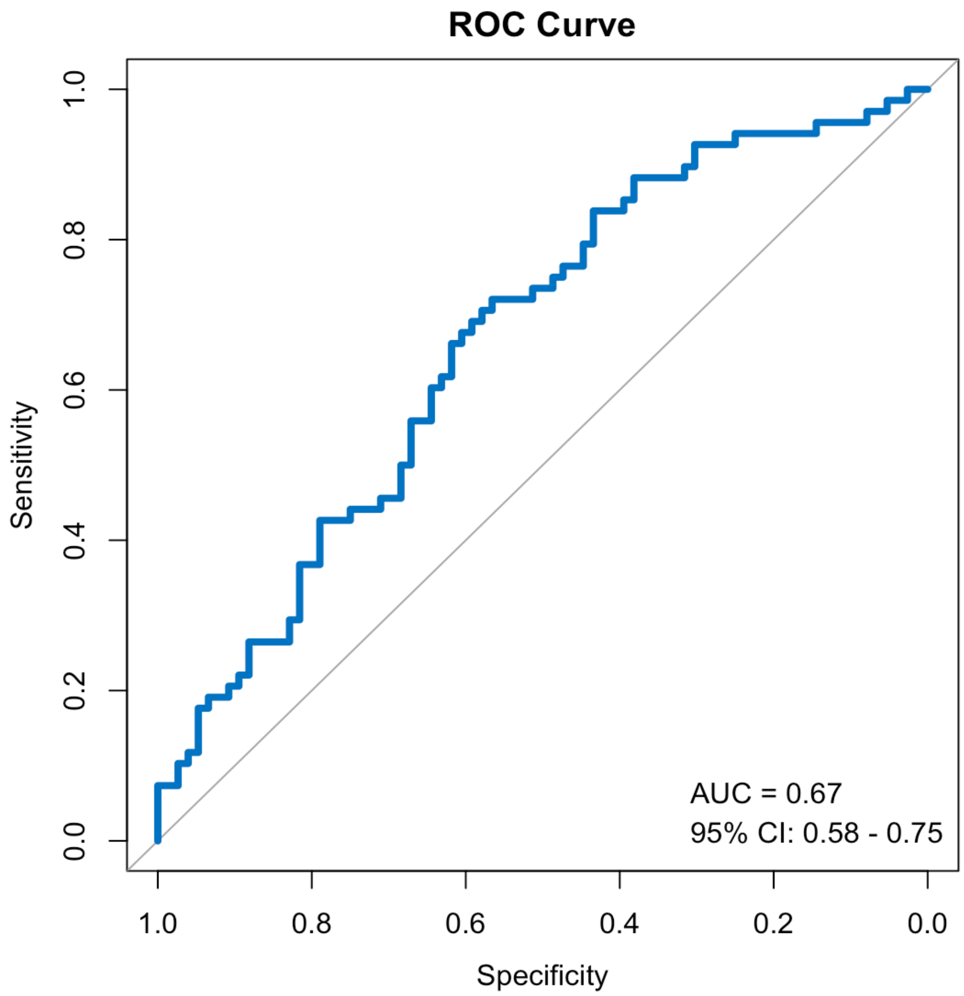


*Supplementary Fig. S3.* Predicting whether a participant has a clinical diagnosis or not using effective connectivity parameters of the basal forebrain. The receiver operating characteristic (ROC) curve depicts the results of the parametric empirical bayes leave-one-out cross-validation procedure, demonstrates the balance between sensitivity and specificity across various thresholds. The area under the curve (AUC) quantifies the model's capacity to accurately classify a new participant as being a clinical participant or not.

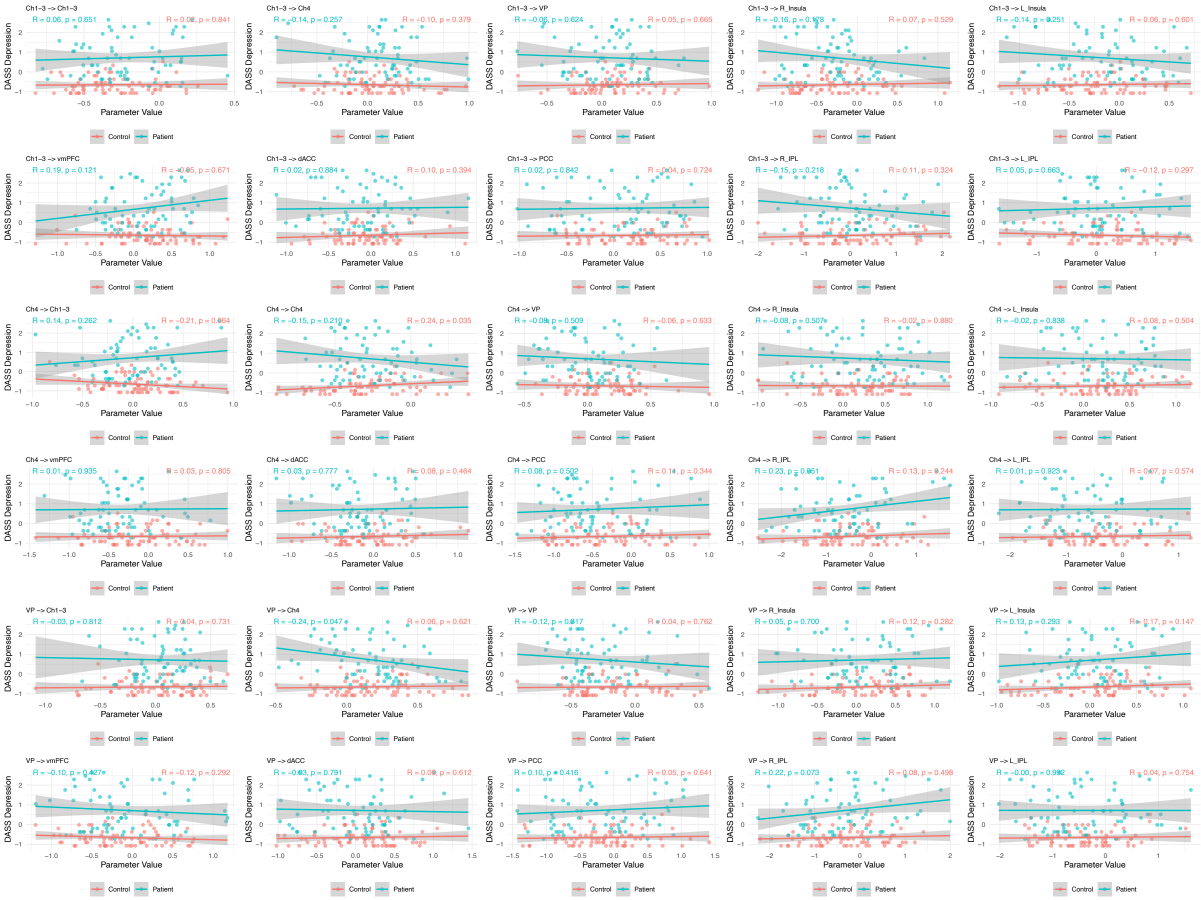

*Supplementary Fig. S4.* Scatterplot and correlations between the expected values for connectivity parameters originating in basal forebrain subregions and depressive symptom severity (assessed through DASS Depression subscale). Healthy controls are shown in salmon and clinical participants are shown in cyan. Shaded area indicates 95% CI.

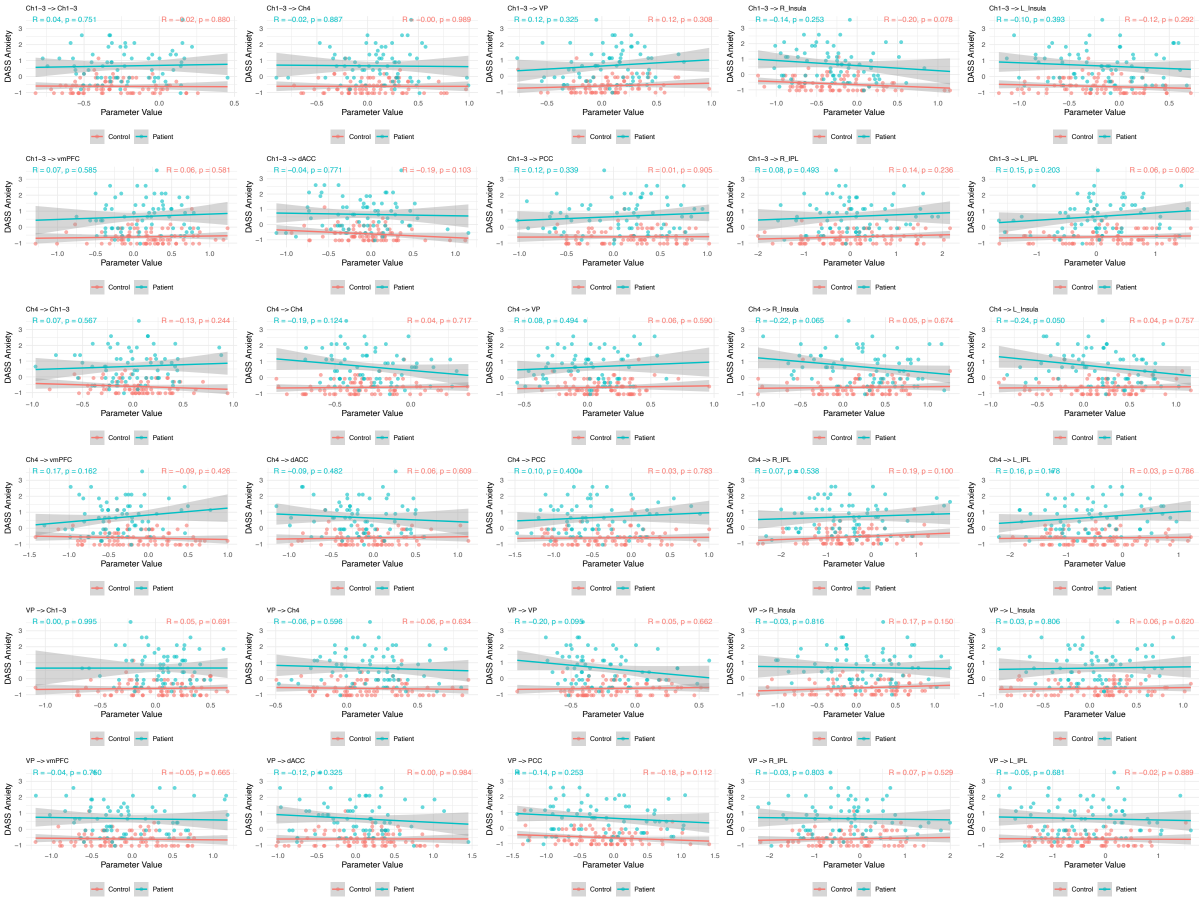

*Supplementary Fig. S5.* Scatterplot and correlations between the expected values for connectivity parameters originating in basal forebrain subregions and anxiety symptom severity (assessed through DASS Anxiety subscale). Healthy controls are shown in salmon and clinical participants are shown in cyan. Shaded area indicates 95% CI.
